# Electrostatics facilitate mid-air host attachment in parasitic jumping nematodes

**DOI:** 10.1101/2025.02.20.639345

**Authors:** Ranjiangshang Ran, Justin C. Burton, Sunny Kumar, Saad Bhamla, Adler R. Dillman, Victor M. Ortega-Jimenez

## Abstract

Jumping can be hazardous for entomopathogenic nematodes (EPNs) as those that fail to attach to an insect host face death by predation or starvation. Recently, it has been shown that electrostatic charges on large insects can prompt a close-range detachment of free-living nematodes, which are non-parasitic and unable to jump. However, it remains unclear if static electricity can influence aerial interactions between parasitic jumping worms and their insect hosts. Here we analyze and model the trajectories of jumping EPNs in still air as they approach fruit flies with varying electrostatic charge. We discover that the nematodes’ attachment to the host is facilitated by an electrical potential of a few hundred volts, a magnitude commonly found in flying insects. A model combining electrostatics, aerodynamics, and Bayesian inference indicates that the electrostatic charge on jumping nematodes is ∼ 0.1 pC, which aligns with theoretical predictions for electrostatic induction. Drag coefficients based on host-nematode interactions in the presence of horizontal wind show differences at both low and high jumping velocities. Numerical simulations show that intermediate wind speeds (∼ 0.2 m/s) can further increase the likelihood of host attachment, as wind-driven aerial drifting allows the worms to reach hosts at greater distances. Our results suggest that submillimeter parasites that become airborne may exploit the electric charge carried by their host to facilitate attachment and thus enhance survival. The use of quantitative physical models provides valuable insights into understanding complex airborne infectious diseases mediated by natural environmental forces.

**Significance Statement:** Entomopathogenic nematodes (EPNs) are submillimeter parasites renowned for their explosive aerial jumping, allowing them to reach distant insect hosts. They serve as important model organisms and natural biopesticides. Our work reveals that these tiny organisms can be electrostatically attracted to charged hosts, such as fruit flies, increasing the likelihood of infection. Experiments show that host attachment is significantly enhanced by electrostatic forces generated by naturally occurring electric fields from flying insect hosts. Our computational model confirms that the static charge of EPNs agrees with theoretical predictions from electrostatic induction. We propose that electrostatics play a crucial role in enhancing the survival of these jumping parasites and provide a framework for modeling environmental forces in aerial parasite-host interactions.

---

Static electricity is a common phenomenon in the atmosphere, playing a fundamental role in the ecology of small organisms, such as terrestrial invertebrates (1), as well as affecting interspecific interactions, such as pollination (2–9), predation (10–12), and parasitism (13, 14). For example, key pollinators such as bees (4, 6), lepidopterans (7), hoverflies (8), and hummingbirds (9) often accumulate positive electrostatic charges via triboelectric effects, while flowers and pollen usually carry negative charges (5, 15, 16). This potential difference results in electrostatic forces strong enough to induce contactless pollen transfer, thus enhancing pollination (1, 2, 5). In predator-prey relationships, it has been shown that the silk of spider webs can be electrostatically attracted to and deformed by positively charged insects (10), facilitating prey capture (11). Interestingly, ballooning spiders can take advantage of electric fields in the atmosphere and drift through the electrified sky on their silk strands over large distances (17, 18). Electrostatics also influence antipredatory mechanisms and parasitism. Caterpillars can detect the electric field emitted by their electrostatically charged predators to avoid them (12). Hummingbird flower mites sense electric field from hummingbirds, allowing them to hitch rides and colonize new flowers (19). Even small, slow-moving parasites such as ticks can be electrostatic attracted to their animal hosts, increasing their likelihood of attachment (13, 14). Despite these significant advances in understanding electrostatics in organismal ecology, the physics underlying these complex biological interactions remain unclear, particularly in aerial interactions within mesoscale ecological systems.

Entomopathogenic nematodes (EPNs) are unique among roundworms due to their extraordinary abilities to jump into the air, reaching heights up to 25 times their size (20), enabling them to reach their insect hosts. They have been studied as both model organisms and biopesticides against insect pests (21, 22), as well as for their relationship with symbiotic bacteria that they utilize for infection (23–25). In particular, the jumping abilities of the EPNs *Steinernema carpocapsae* at the infective juvenile stage have been previously described (20, 26–29). Since *S. carpocapsae* do not feed until they reach a host at this stage, it is crucial for them to successfully land on their host after jumping, otherwise they face high risks of predation, desiccation, and starvation (30–32). Recent evidence suggests that the free-living nematodes *Caenorhabditis elegans*, which are non-parasitic and unable to jump, can be detached by electrostatically charged bumblebees, potentially enhancing their dispersal (33). However, it remains both experimentally and theoretically unclear whether the successful host attachment of jumping nematodes is affected by electrostatics forces and the presence of wind. Furthermore, the charging mechanism of jumping EPNs—whether it is bioelectricity, triboelectrification, or electrostatic induction—remains a mystery.

Here we investigate the effects of electrostatics on EPNs-host interactions in mid-air. We analyze and model the trajectories of these jumping worms during takeoff and landing in the presence of fruit flies with varied electrical potentials. We find that the nematode’s successful attachment increases with the fly’s electric potential. Drifting was observed in nematodes jumping through the airflow generated by a horizontal wind tunnel. A theoretical model integrating electrostatics, aerodynamics, and Bayesian inference reveals that the electrostatic charge on the nematodes is ∼ 0.1 pC, consistent with predictions from electrostatic induction. Our inference method determines key aerodynamic properties of EPNs, such as hydrodynamic radius and drag coefficient, which we then use to construct a fully quantitative model of host-parasite interactions. Moreover, numerical simulations reveal that while increasing electric potential monotonically enhances host attachment, intermediate wind speeds (∼ 0.2 m/s) further improve attachment by allowing the worms to reach more distant hosts. Overall, our results provide a framework for developing physics-based models of host-parasite interactions that can be generalized to other mesoscale ecological systems.

## Results

### Electrostatic attraction of jumping EPNs

Infective juveniles of *S. carpocapsae* were used to test the electrostatic effects produced from a fruit fly, *Drosophila melanogaster*, with varied electrical potentials. The nematodes are ≈ 25 µm in diameter and ≈ 400 µm long (20). A drop of water containing active nematodes were deposited onto a sheet of folded wet filter paper (see *SI* Fig. S1). This wet paper was mounted on a metal stand connected to the ground. A living fruit fly was tethered to a copper wire and connected to a high-voltage power supply (see *Materials and Methods*). The tethered fly was positioned at two different heights (5.1 mm and 6.2 mm) above the grounded wet paper. Flying insects such as bumblebees (4), honeybees (34, 35), and houseflies (36) are reported to naturally accumulate static charge of 10−200 pC through triboelectric effects, corresponding to voltages of 50−1000 V (10, 37). Thus, the voltage on the tethered fly was adjusted from 100−700 V relative to the ground. Nematodes that successfully stood and jumped (20) were recorded using a high-speed camera (see *Materials and Methods*).

We also conducted similar experiments using a charged metal sphere with a diameter of 2.54 mm instead of an insect. Our results confirm that both the charged fly and the metal sphere attract jumping worms in mid-air. The takeoff, aerial phase, and attachment to the charged host of a jumping nematode are shown in Fig. 1*A*. The jumping process, including loop formation and jumping, is shown in Fig. 1*B* and was discussed in detail in a previous work (20). We note that after launch, and despite that some individual presented an initial heading in the opposite direction, the rotating worms were pulled toward and landed on the insect host. As shown by the tracked trajectories of multiple worms in Fig. 1*C*, nematodes took off at a maximum jumping speed of *U*_0_ ≈ 1.5 m/s, followed by a period of deceleration due to aerodynamic drag and gravity force. Therefore, these experiments confirm that electrostatic charges carried by tiny insects, such as a fruit fly, are enough to induce mid-air attraction and facilitate attachment of jumping nematodes to the host.

**Fig 1.**
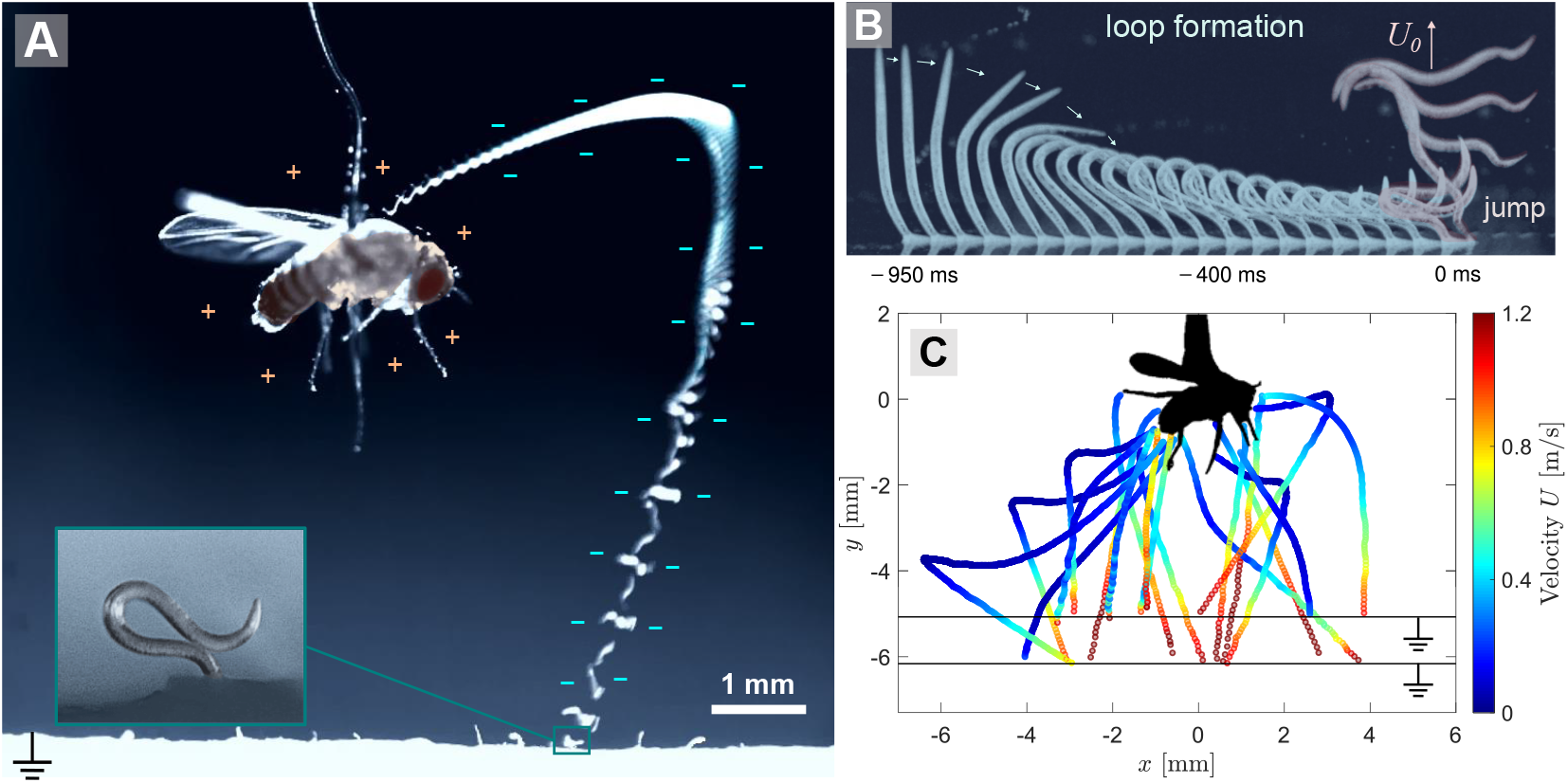
Jumping nematodes electrostatically attracted to a charged insect host. (*A*) Trajectory and body orientation of a jumping nematode pulled towards a positively charged fruit fly. Inset: a zoomed in color photograph of a nematode in looping formation on a wet paper. (*B*) Sequential snapshots of a *Steinernema carpocapsae* nematode’s jumping performance: loop formation from –950 ms to 0 ms and final unleashing and takeoff within 0 ms to 5 ms. (*C*) Trajectories of jumping nematodes’ center of mass (*N* = 19) colored by their instantaneous velocity, *U*. In these experiments, the nematodes took off from a grounded plane at two different heights, roughly 5.1 mm and 6.2 mm below the fruit fly, respectively. The voltage on the charged fruit fly was varied from 100 V to 700 V.

### Electrostatic model of host-parasite interaction

To further understand the physical principles behind the trajectories of jumping EPNs as they approach the charged host, we developed a theoretical model of host-parasite interactions. The insect host was modeled as an isolated sphere with a positive charge *Q* = 4*πϵ*_0_*a*_*f*_ *ϕ*, where *ϵ*_0_ = 8.854 × 10^*−*12^ F/m is the vacuum permittivity, *a*_*f*_ = 1 mm is the equivalent radius of the fruit fly in Fig. 1*A*, and *ϕ* is the host’s electric potential with respect to the ground. The nematodes were also modeled as spheres with a negative charge −*q*, whose values were determined from trajectory fitting. The grounded plane where the nematodes took off was modeled by placing a negative image charge −*Q* symmetrically below the plane (38). The image charge of the nematode was neglected since it is only important very close to the plane. The model and the associated electric field are illustrated in Fig. 2*A*. We have made two physical assumptions to simplify computations, i.e., the host is modeled as an isolated spherical capacitor, and the wet filter paper is modeled as a grounded plane. We justify the use of these assumptions by quantifying the error they introduce in the *SI* (39).

**Fig 2.**
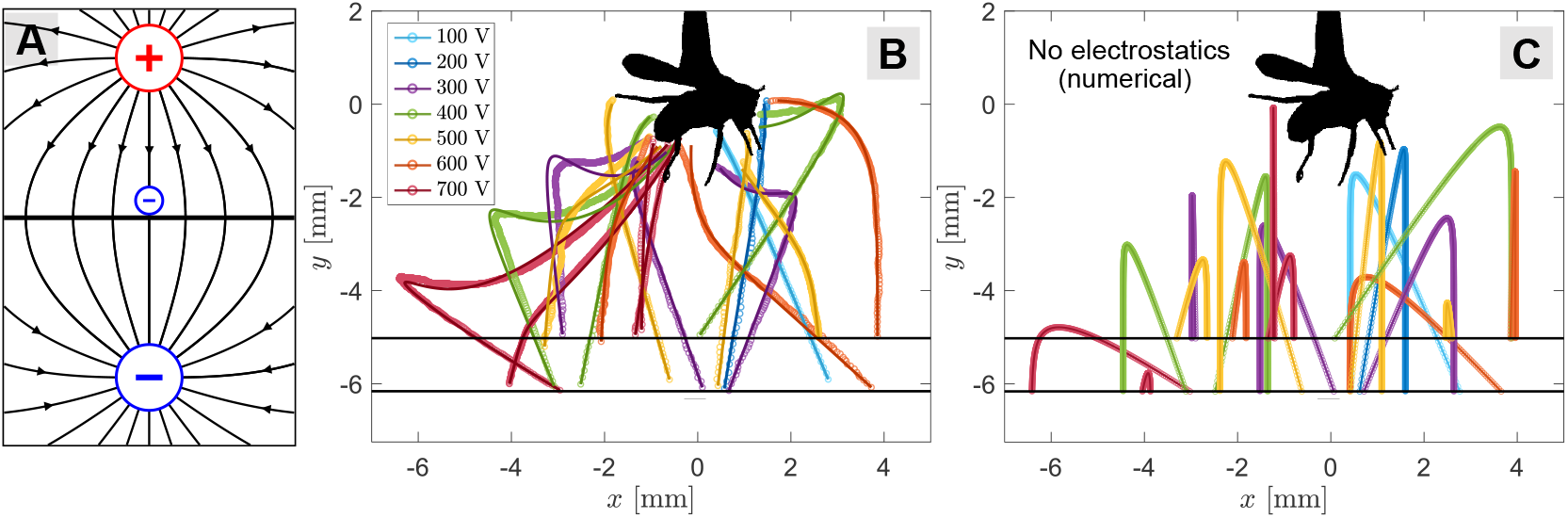
Electrostatic model and trajectory fitting of jumping nematodes. (*A*) The fruit fly is modeled as a sphere with a uniform positive charge +*Q*, the jumping nematode is modeled as a smaller sphere with a negative charge −*q*, and the grounded conducting plane is replaced by a negative image charge −*Q* located symmetrically below the plane. The electric field is illustrated by black lines with arrows. (*B*) 2D projection of the 3D model fitting results using the nematodes’ trajectories in Fig. 1*C*. Circle symbols are experimental data and solid curves are fitting results using Eq. (1). Different colors represent different voltages on the fruit fly, ranging from *ϕ* = 100-700 V. (*C*) Numerical integration of the same nematodes’ trajectories as in Fig. 2*B*, but this time with electrostatic effects removed by setting *q* = 0. Colors in this plot represent trajectories that share the same initial condition in Fig. 2*B*. Only 1 out of the 19 trajectories land on target without electrostatics.

The equation of motion of the jumping nematodes with this simplified model is given by:

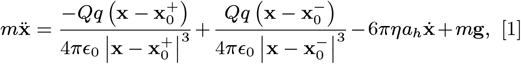

where *m* and **x** are the mass and the position of a nematode, respectively, 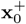 is the position of the positively charged host, and 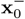 is the position of the negative image charge. The first and second terms on the right-hand side (RHS) of Eq. (1) represent the electrostatic attraction and repulsion from the charged host and the image charge, respectively. Besides electrostatic forces, the nematode also experiences aerodynamic drag force and gravity, which are modeled by the third and fourth terms on the RHS of Eq. (1), where *η* = 1.849 × 10^*−*5^ Pa.s is the dynamic viscosity of ambient air, *a*_*h*_ is the hydrodynamic radius of the nematode, and **g** = −*g***ŷ** is the gravitational acceleration with *g* = 9.81 m/s^2^. We note that the drag force is modeled by Stokes law (40), but the Reynolds number of a nematode can be greater than unity at high jumping velocities. Thus, *a*_*h*_ is taken as a fitting parameter and varies with each nematode.

The trajectories of jumping nematodes can be highly three-dimensional (3D), which is evidenced by the blurred images of their bodies resulting from their movement in and out of the camera’s focal plane (see Movie S1). Since electrostatic forces depend on the 3D distance between the parasite and the host, it was necessary to fit the experimental trajectory data to the theoretical model in Eq. (1) in three dimensions. However, we only captured the two-dimensional (2D) trajectories of the nematodes with one camera. To tackle the missing observables (out-of-plane displacement and velocity), we resorted to a Bayesian inference method known as Markov Chain Monte Carlo (MCMC), which allowed us to reconstruct the nematodes’ 3D trajectories with 2D data and simultaneously infer their charge *q* (see *Materials and Methods*). Figure 2*B* shows the 2D projection of the 3D fitting results for the nematodes’ trajectories (full 3D trajectories are shown in *SI* Fig. S3). We find that the model in Eq. (1) accurately captures the nematodes’ trajectories, especially the sharp turns caused by electrostatic forces. We also fitted the trajectories in analogous experiments with a charged metal ball instead of a fruit fly, and found that our model fit the experimental data equally well (see *SI* Fig. S4). This validates our assumption of treating the irregularly shaped fruit fly as a charged sphere.

To test the role of electrostatics in host attachment, we performed numerical simulations of the trajectories of jumping nematodes using the same initial conditions and fitted parameters as in Fig. 2*B*, but this time with electrostatic effects removed by setting their charge *q* to zero. Figure 2*C* shows the numerical results of the hypothetical nematodes’ trajectories. Without electrostatic effects, nematodes continue in their initial jumping direction and fall straight downward at a terminal velocity due to aerodynamic drag; even those nematodes that jump directly toward the host may fail to reach it due to insufficient jumping velocity. Only 1 out of 19 trajectories successfully reached the insect host. In contrast, when electrostatics are present, all 19 nematodes successfully landed on their target host (Fig. 2*B*). These results suggest that electrostatic effects play a fundamental role for the successful attachment of nematodes to insects.

### Charging mechanism in EPNs

How are jumping nematodes charged? Two possible charging mechanisms include electrostatic induction and triboelectrification. We hypothesize that induction is the most likely mechanism, as nematodes were consistently attracted to the host in our experiments. It is important to note that the relative humidity in the experimental setup was near saturation and the paper was wet. Furthermore, it was recently shown that nematodes have a water coating on the surface of the body, which allows the formation of a capillary latch during looping formation (20). Thus, in our model the worms were considered as conductors in contact with the grounded plane, making induction possible. Charging via induction can be visualized in 3 steps, as illustrated in Fig. 3*A*. Initially, the nematode undergoes polarization when a nearby host with a positive charge causes the mobile charges within the nematode to separate. Contact with a grounded plane then results in an electric current of positive charge to the ground (or, electrons moving from the ground to the nematode). Finally, during detachment, the nematode, now carrying a negative charge, jumps away from the ground and is attracted to its positively charged host.

**Fig 3.**
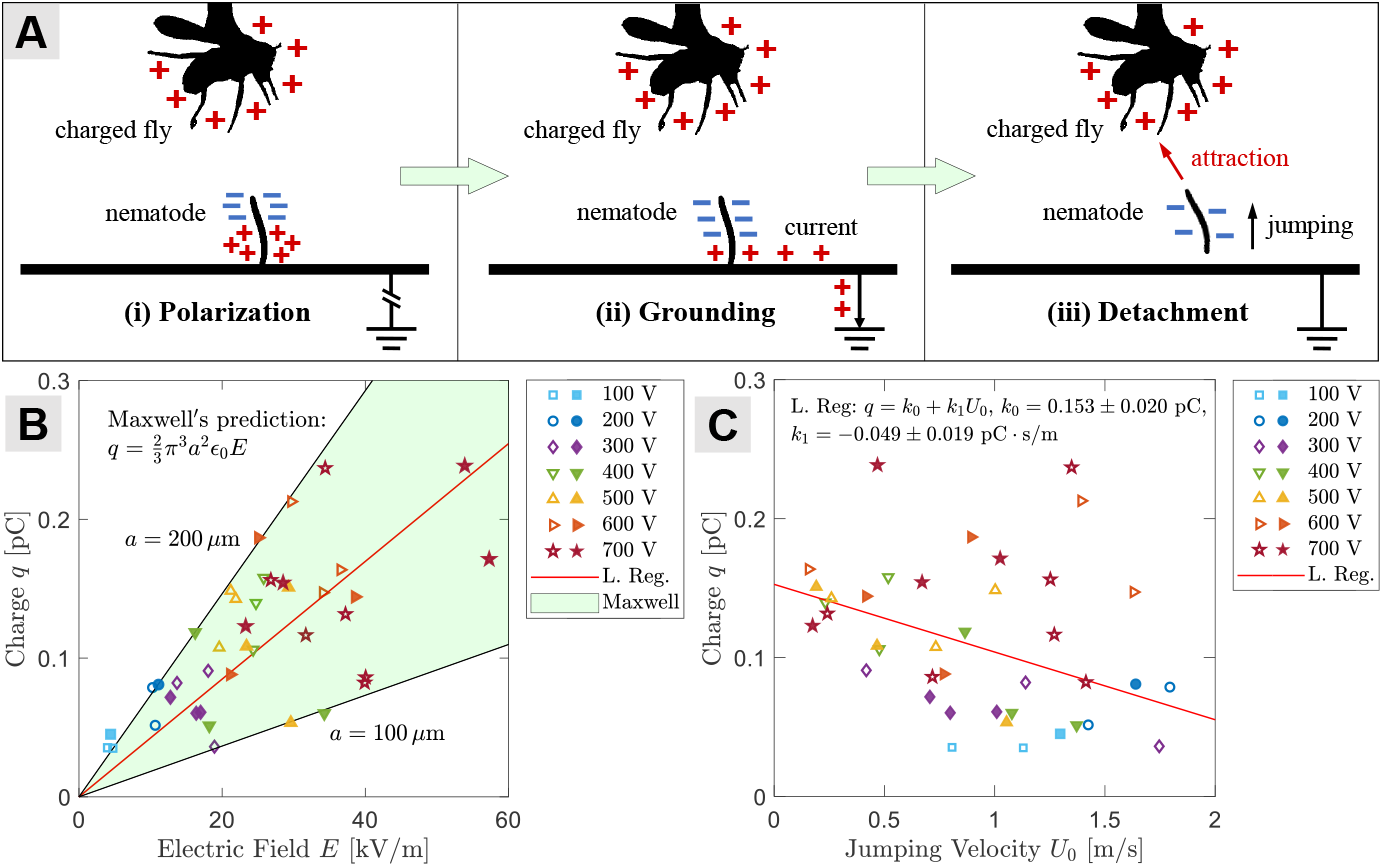
Nematode charging mechanism and Maxwell’s predictions. (*A*) Illustrations of the induction charging mechanism in jumping nematodes. (i) Polarization: the positive charge on the host nearby causes the mobile charges in the nematode to separate. (ii) Grounding: when in contact with a grounded plane, an electric current arises, positive charge leaves the nematode and enters the ground (e.g., electrons move from the ground to the nematode). (iii) Detachment: as the jumping nematode detaches itself from the ground, it carries the opposite (negative) charge and is attracted to its host. (*B*) The inferred charge on the jumping nematodes obtained from trajectory fitting, *q*, as a function of the electric field magnitude at the location where they jump, *E*. The shaded area represents Maxwell’s prediction for the charge on a conducting sphere sitting on a conductive plane: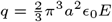, with a lower bound on the sphere’s radius of *a* = 100 µm and an upper bound of *a* = 200 µm. The solid red line is the linear regression of the data, resulting in *a* = 154 µm. (*C*) The inferred charge on the jumping nematodes, *q*, as a function of their jumping velocity, *U*_0_. Nematodes with a lower charge require a larger jumping velocity to land on target. The solid red line represents the linear regression: *q* = *k*_0_ + *k*_1_ *U*_0_, with the values and standard errors of *k*_0_ and *k*_1_ reported in the graph. In (*B*) and (*C*), solid symbols are experimental data with a charged fruit fly, and open symbols are experimental data with a charged metal sphere.

To confirm the induction mechanism, we compared the nematodes’ charge from Bayesian inference, *q*, to theoretical predictions based on the electric field magnitude, *E*. Since *q* remained a constant during flight after the nematodes detached from the grounded plane, *E* was calculated at the location where the nematodes jumped, **x**_0_:

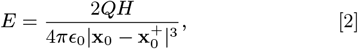

where *H* denotes vertical distance between the nematodes and the host. Figure 3*B* shows *q* as a function of *E*. We find that *q* ranges from 0.05 to 0.25 pC, increasing with the electric field magnitude (*E*) and the host’s electric potential (*ϕ*). Theoretically, the charge on a conducting sphere of radius *a* in contact with a conducting plane of a known surface charge density *σ* was first solved by Maxwell (38, 41):

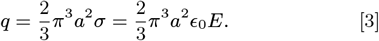

Remarkably, we find that all *q* data are bounded by the shaded area in Fig. 3*B*, which represents Maxwell’s prediction [Eq. (3)] for *a* = 100 – 200 µm. The linear regression of all data (red line) yields *a* ≈ 154 µm. Although the actual shape of the worm is cylindrical, and we treat the worms as spherical, this range of length scales is in excellent agreement with the typical worm size (Fig. 1*A*), validating our hypothesis that charging is achieved through electrostatic induction. Moreover, our numerical simulations show that the charge on conducting upright cylinders via induction is also of order 0.1 pC (see *SI* Fig. S8), which agrees with the inferred charge on nematodes and Eq. (3).

Despite the excellent agreement with theoretical predictions for electrostatic induction, triboelectric effects could still play a role at high jumping velocities (e.g., *U*_0_ ≥ 1.5 m/s). If this is the case, then one might expect *q* to increase with *U*_0_. However, we find an opposite trend, that *q* is negatively correlated with *U*_0_, as shown in Fig. 3*C*. A linear regression, *q* = *k*0 + *k*_1_*U*_0_, confirms the negative correlation with *k*_1_ = − 0.049±0.019 pC · s*/*m (reported with standard error). The negative correlation is further supported by an *F*-test using analysis of variance (42), showing a *F*-statistic of 6.48 and a *p*-value of 0.015, indicating that linear regression fits significantly better than a degenerate model with only a constant term, at the 5% significance level (*p <* 0.05). The negative correlation is likely due to selection bias in our data: we only analyzed experimental trajectories where nematodes eventually attached to the host, and nematodes with less charge generally required larger jumping velocities to reach their host target. More simply, Fig. 3*C* reflects the fact that electrostatics facilitate host attachment by decreasing the necessary jumping velocity of parasitic nematodes.

### EPNs drifting in the wind and electrostatics

Wind is ubiquitous in nature and can influence electrostatic attachment in host-parasite interactions. To investigate aerial drifting in jumping EPNs due to horizontal winds, we filmed nematodes jumping on a flat paper surface under airflow conditions generated by a wind tunnel (see *Materials and Methods*). The trajectory and body orientation of a nematode drifting in the wind are shown in Fig. 4*A*). The wind speed was set to *U*_*∞*_ ≈ 0.2 m/s. At this speed, the boundary layer over a flat plate is typically laminar, and can be modeled by the classical Blasius solution (see *Materials and Methods*). Figure 4*B* shows the velocity profile and boundary layer thickness from the Blasius solution, along with the trajectories of the nematodes’ center of mass and model fitting results using Bayesian inference. Once again, our aerodynamic model accurately captures the experimental trajectories of the nematodes. Notably, we find that most individuals were able to reach heights over 5 mm (≈ 12 times their length), allowing them to intersect the edge of the 99% velocity boundary layer (i.e., the height where the wind speed is 0.99 *U*_*∞*_). As a result, the nematodes are able to access and be carried away by the free stream outside the boundary layer, facilitating long-range dispersal.

**Fig 4.**
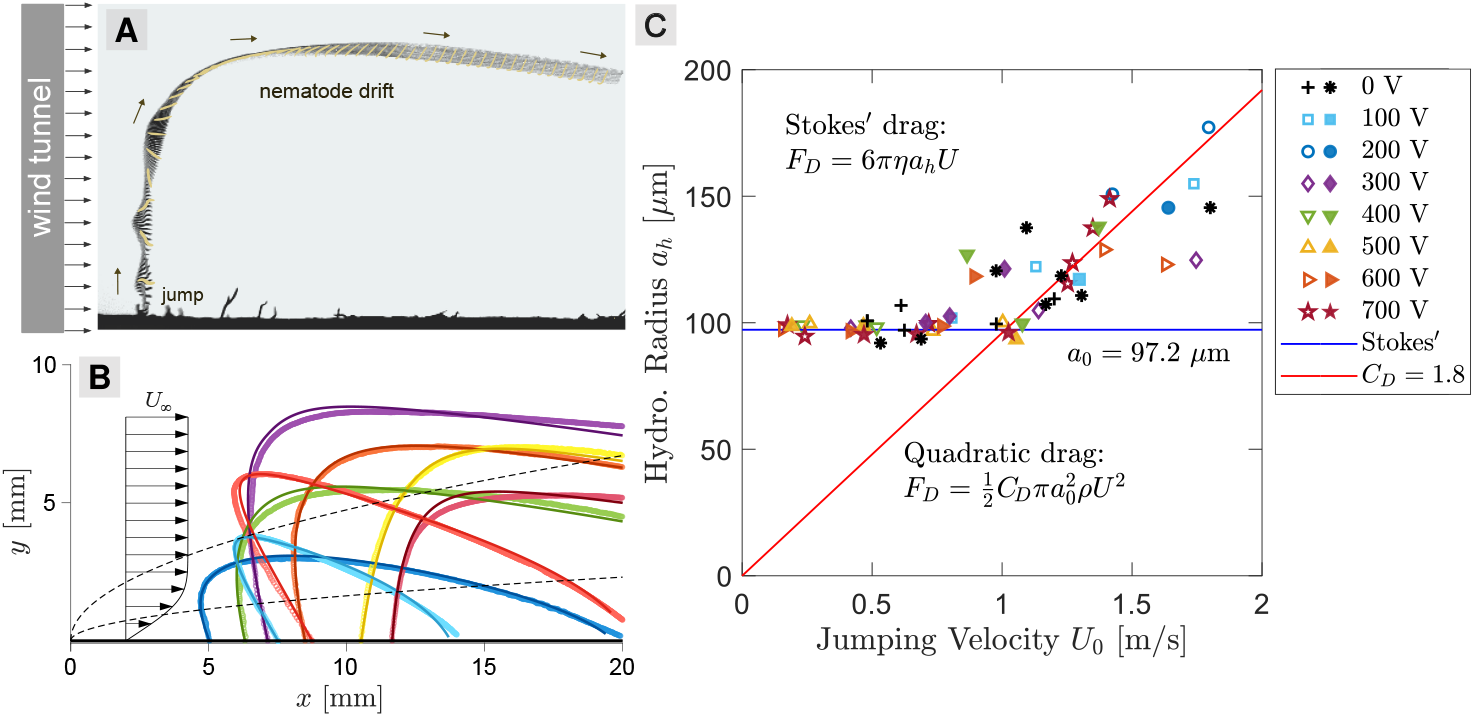
Aerial drifting of jumping nematodes in wind. (*A*) Trajectory and body orientation of a jumping nematode drifting in a horizontal laminar flow without electrostatic effects. (*B*) Trajectories of the center of mass of jumping nematodes (*N* = 8) in wind’s boundary layer near a flat plate. Circle symbols are experimental data; solid curves are model fitting results using the Blasius solution of a laminar boundary layer in Eq. (8). Black arrows illustrate the velocity profile in the free stream and in the Blasius boundary layer. Black dashed curves represent the 99% velocity boundary layer thickness (top) and the displacement boundary layer thickness (bottom). (*C*) The inferred hydrodynamic radius of the jumping nematodes, *a*_*h*_, as a function of their jumping velocity, *U*_0_, for experiments both with and without electrostatics. For electrostatic experiments (100 – 700 V), solid symbols are experimental data with a charged fruit fly, and open symbols are experimental data with the fly replaced by a charged metal sphere. For data without electrostatics (0 V), asterisk symbols (*) are experiments with nematodes jumping in wind, and plus symbols (+) are control experiments with nematodes jumping in still air. At low velocities (*U*_0_ ≲ 1 m*/*s), the drag force on the nematodes follows Stokes’ law: *F*_*D*_ = 6*πηa*_*h*_*U*, with an average hydrodynamic radius of *a*_0_ ≈ 100 µm (solid blue line). At high velocities (*U*_0_ ≳ 1 m*/*s), the hydrodynamic radius increases with *U*_0_ due to the transition from the linear Stokes’ drag to a quadratic relation: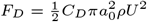, with an estimated drag coefficient of *C*_*D*_ ≈ 1.8 (solid red line).

Our inference method estimates the hydrodynamic radius, *a*_*h*_, of jumping nematodes in electrostatic experiments (i.e., jumping and attracted by a charged fly or a metal ball), and wind experiments (i.e., jumping in a horizontal wind and in still air). Figure 4*C* shows *a*_*h*_ as a function of the jumping velocity *U*_0_. The consistent inference of *a*_*h*_ between wind and electrostatic experiments is remarkable, and indicates that our method independently infers the aerodynamic and electrostatic properties of the nematodes. At lower jumping velocities (*U*_0_ ≲ 1 m*/*s), we find that *a*_*h*_ remains nearly constant, with an average of *a*_0_ ≈ 100 µm. This value lies in between the nematodes’ diameter (25 µm) and body length (400 µm), and is an average of their spinning body during flight. At higher jumping velocity (*U*_0_ ≳ 1 m*/*s), we find that *a*_*h*_ increases with *U*_0_. We believe that this is because the Stokes drag used in our model, *F*_*D*_ = 6*πηa*_*h*_*U*, is only valid at low Reynolds numbers. The Reynolds number of the EPNs is defined as: Re = *ρU*_0_*a*_*h*_*/η*, with *ρ* being the air density. For *U*_0_ = 1 m/s and *a*_*h*_ = 100 µm, the Reynolds number is Re ∼ ℴ (10^1^). Therefore, we expect a transition to a quadratic relation between the velocity and the drag force, 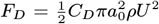. Indeed, a linear regression of the high *U*_0_ data yields a drag coefficient of *C*_*D*_ = 1.8, which is also an average over the spinning motion of the nematode.

With quantitative information of the electrostatic and aerodynamic properties of jumping EPNs, we performed numerical simulations of nematodes drifting in wind with the addition of a charged host nearby (see *SI* and Movie S4). In our simulations, the nematodes were launched from random positions and takeoff angles at *U*_0_ = 1 m/s, and “capture” was defined as successful host attachment. Figure 5 shows the probability of capture, *P*_*c*_, a function of the wind speed (*U*_*∞*_) and the host’s electric potential (*ϕ*). We find that *P*_*c*_ increases monotonically with *ϕ* for all *U*_*∞*_ values. In still air (*U*_*∞*_ = 0), *P*_*c*_ increases from 10% at *ϕ* = 100 V to more than 60% at *ϕ* = 700 V. Our results indicate that electrostatics consistently facilitate host attachment in jumping nematodes. Moreover, we find that intermediate wind speeds (*U*_*∞*_ ≈ 0.2 m/s) can further increase the likelihood of host attachment, especially at higher *ϕ*. At *ϕ* = 800 V, intermediate wind speeds increase *P*_*c*_ from 60% to more than 70%. Thus, aerial drifting allows nematodes to be electrostatically attracted to more distant hosts downstream (see Movie S5).

**Fig 5.**
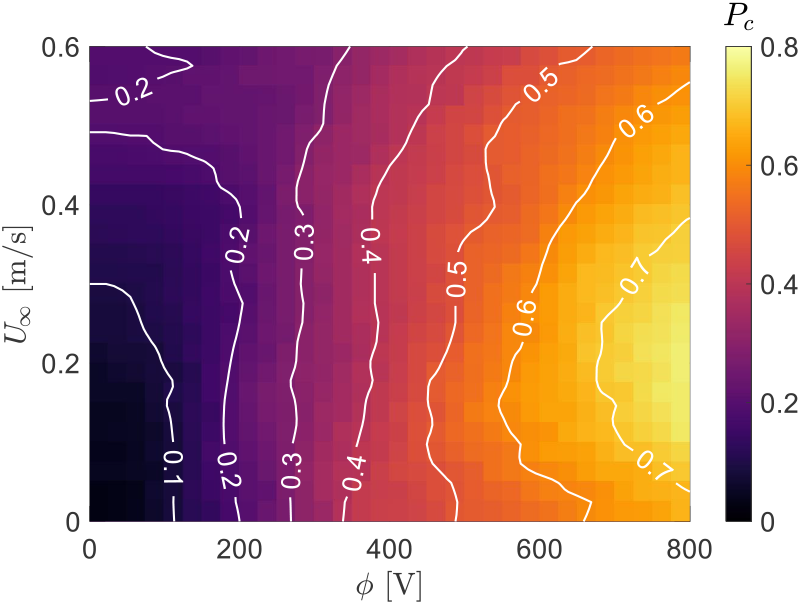
Capture probability of jumping nematodes (*P*_*c*_) in an assortment of numerical simulations varying the wind speed (*U*_*∞*_) and the host’s electric potential (*ϕ*). While *P*_*c*_ increases monotonically with *ϕ*, intermediate wind speeds (*U*_*∞*_ ≈ 0.2 m*/*s) lead to a higher probability of capturing, especially at larger *ϕ*.

## Discussion and Summary

EPNs are submillimeter-sized parasites that prey on insects and are unique in the phylum Nematoda for their ability to propel themselves into the air, traveling up to 25 times their size (20). It has been suggested that these jumping skills in roundworms can facilitate host seeking and attachment (26–28). Here, we demonstrated through experiments and theoretical modeling that electrostatic forces significantly increase the likelihood of nematode attachment to their charged host in mid-air. Thus, jumping nematodes do not need to precisely predict their jump to land on a highly mobile target; they only require to get close enough for electrostatic forces to induce attraction and ensure successful infection. Individual insects can become easily charged while moving on surfaces, in the air, or through direct contact with charged objects, and carry tens to a thousand picocoulombs (1), corresponding to electrical potentials of several hundred volts (10). Moreover, a recent report suggests that insect swarms, such as those of honey bees, can generate local charge densities comparable to those produced by thunderstorms (37). It is important to notice that we used small insects (fruit fly size ≈ 3 mm) as hosts to evaluate electrostatic effects on EPNs. However, we anticipate that larger insects, such as craneflies, beetles, or bees, which carry larger charges, could further increase the likelihood of nematode attachment, especially when these insects are moving in swarms. Moreover, it is expected that charged insects moving closer to nictating nematodes will also have an increased chance of infection, even without the nematodes actively jumping.

We observed that a plastic syringe rubbed on human hair, as well as a charged water droplet, placed near a standing nematode (Movie S6) or a group of nematodes (Movie S7), produced worm detachment and attraction. This agrees with findings on the electrostatic pulling of *C. elegans* (free-living, non-parasitic nematodes that do not jump) by charged bodies (7). Other organisms such as grounded ballooning spiders, seem capable of harnessing atmospheric electricity to passively ascend into the sky (17). This possibility of aerial dispersal via atmospheric electrostatics in EPNs is intriguing. Given their need for water to survive, an electrostatic ascent could facilitate interaction with tiny drops in clouds, allowing them to fall back to the ground as rain, which may explain their global distribution. Future studies exploring how atmospheric electrostatics influence jumping nematodes is necessary to better understand their dispersal capabilities.

We showed that EPNs are attracted to charged hosts through electrostatic induction, which is a common charging mechanism among tiny organisms and biological materials (1, 10, 11, 14, 33). It does not require direct contact or rubbing with the charged body and can occur at a relatively large distance compared to the body length. Additionally, a body with either positive or negative charge can produce attraction on a grounded biological entity, such as a tick (13) or a spider web (10, 11). Our results indicate that jumping roundworms are charged ∼ 0.1 pC due to charge separation induced by a positively charged host, which agrees with findings on the charging mechanisms of free-living nematode *C. elegans* (33). Consequently, we expect that negative charges will elicit similar attraction responses on jumping EPNs. Research on the role of the water coating on nematode bodies is needed to understand how it affects charge mobility and induction.

Wind is a major driver of dispersal for small organisms, ranging from bacteria to spores to wingless arthropods to plant seeds. For example, pollen grains and dandelion seeds can be carried by the wind over distances of several kilometers, depending on environmental conditions (43, 44). The diameter of pollen grains is similar to the body thickness of nematodes, suggesting that wind could also carry jumping nematodes over similar kilometer-scale distances. In unfavorable environmental conditions and with low prey availability, wind dispersal may enhance the chances of nematode survival. Our numerical simulations support this, showing that intermediate wind speeds (∼ 0.2 m/s) increase the likelihood of worms reaching and attaching to an electrostatically charged host. Field and lab research is required to better understand the long-distance dispersal of nematodes through wind and electrostatics.

Our findings were only made possible by quantitative modeling of the 3D jumping trajectories of nematodes using Bayesian inference. The inference method was crucial; the model requires simultaneous inference of physical parameters such as electrostatic charge and hydrodynamic radius in addition to the out-of-plane velocity that is missing from the 2D images. Importantly, the inferred magnitude of charge on the EPNs revealed induction as the primary electrostatic mechanism at play, and showed excellent agreement with Maxwell’s theory. This also suggests that EPNs, and perhaps other mesoscale organisms, can be modeled as conductors when enough water is present (i.e., high relative humidity). Because of this, we expect that our methods can be applied to a variety of biological systems where electrostatics and other natural environmental forces are important, but are yet to be discovered.

To summarize, there are three major takeaways from this study. First and foremost, electrostatics clearly provide a significant enhancement to the capture probability of EPNs by their hosts. A few hundred volts, often encountered in insects, induce an opposite charge and apply an attractive force on the EPNs after jumping. Secondly, the charging mechanism in jumping nematodes, in the presence of a distant charged host, is driven by electrostatic induction. Third, wind also increases the probability of host attachment in nematodes, particularly when they drift over long distances and specially if the host is electrostatically charged. In fact, the probability of capture without electrostatics was remarkably low, and may suggest electrostatic forces are necessary for jumping to be a successful host attachment strategy. Finally, our results show that electrostatic forces play a fundamental role in enhancing nematode attachment to their highly mobile insect hosts, thereby promoting successful infection. We propose that our numerical inference method can be applied to investigate the effects of environmental factors such as electrostatic and aerodynamic forces on aerial host-parasitic interactions.

## Materials and Methods

### Electrostatic effects on jumping EPNs

Infectious juveniles of *Steinernema carpocapsae* were cultured from infected waxworms using white traps (28). A drop containing a multitude of active worms was placed on the upper edge of a filter paper, which was oriented vertically (*SI* Fig. S1). To prevent desiccation of the worms, the paper was kept moist and mounted on a grounded metal stand. Four glass microscopic slides were glued together to form a clear chamber. Inside the chamber, we tethered a fruit fly to a copper wire, which was connected to a high-voltage power supply (ES5P-10W, Gamma High Voltage Research, Inc.). Voltages ranging from 100 V to 700 V were tested. Similar experiments were conducted using a charged metal sphere (diameter = 2.54 mm). A high-speed camera (Nova S6, Photron USA, Inc.) was used to film at 10,000 frames/s, capturing only those sequences of jumping worms in still air that were in focus and reached the charged fly or metal sphere.

### Aerial drifting in jumping nematodes

To investigate the effects of the wind on the jumping trajectories of nematodes, we performed experiments using a small wind tunnel. The wind tunnel was built using two computer fans placed at the inlet and a 3D-printed honeycomb grid along with a fine mesh filter at the outlet. Flow speed was controlled using an adjustable power supply by setting the voltage to ≈ 3 volts, and measured with a hotwire anemometer (Koselig Instruments, LLC). The aforementioned wet paper with the nematodes was placed downstream inside a glass chamber (i.e., the test section). We analyzed only the videos of nematodes that were in focus and drifting in the air.

### Markov chain Monte Carlo (MCMC) method

We used the MCMC method employed in a previous work (45) to fit our experimental data to the model in Eq. (1). The model has 6 fitting parameters: **Π** = [*q, a*_*h*_, *u*_0_, *v*_0_, *w*_0_, *z*_0_], where *u*_0_, *v*_0_, *w*_0_ are the components of the nematode’s jumping velocity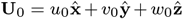, and *z*_0_ is the out-of-plane component of the nematode’s initial position. The nematodes’ mass *m* was taken to be a constant and computed by considering the worm as a cylinder with a diameter of 25 µm, a length of 400 µm, and a water density of 1000 kg/m^3^. Our goal is to find the optimal fitting parameters **Π** that can predict the nematode’s position **X** = [*x, y*] and maximize the conditional probability *P* (**Π**|**X**). Although an explicit expression for *P* (**Π** |**X**) is not available, its values can be computed by Bayes’ theorem:

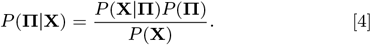

Here, *P* (**X**) = ∫ *P* (**X Π**)*P* (**Π**) *d***Π** is a normalizing constant that can be dropped (46). The probability *P* (**Π**) is assumed to satisfy a multivariate normal distribution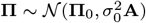, where **Π**_0_ is an initial guess, *σ*_0_ is the uncertainty of the initial guess, and **A** is a diagonal covariance matrix such that each parameter in **Π** is independent. The probability *P* (**X**|**Π**) is defined to minimize the loss function between experimental data and model prediction, ℒ (**X, X**^*′*^) = ∥**X** − **X**^*′*^ ∥, where the double vertical lines denote the *L*^2^-norm. The loss function is assumed to be normally distributed around zero: ℒ (**X, X**^*′*^) ∼ *N* (0, *σ*^2^), and the probability *P* (**X** |**Π**) can be estimated as:

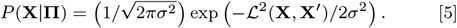

We used the Metropolis-Hastings algorithm (47, 48) to generate a Markov chain of samples from *P* (**Π**|**X**) with the following steps:

Step 0. Propose an initial guess **Π** = **Π**_**0**_.

Step 1. Denote the current state as **Π**. Propose a new state **Π**^***′***^ = **Π** + **A*δ*** using a Gaussian random walk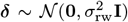, where *σ*_rw_ is the step of the random walk, and **0** and **I** are the zero vector and the identity matrix, respectively.

Step 2. Calculate the acceptance ratio *α*:

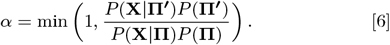

Step 3. Generate a random number *u* from a uniform distribution *u* (0, 1). If *u < α*, accept the new state **Π** = **Π**^***′***^; otherwise reject **Π**^***′***^ and keep **Π** = **Π**. Return to Step 1.

We repeated the above sampling steps for 7,500 to 10,000 times until the loss function converges to its minimum. After the convergence, we continue the sampling for an additional 7,500 to 10,000 times to generate a stationary Markov chain, where all components of **Π** fluctuate around their mean values (see *SI* Fig. S6). We then took the mean values of **Π** on the stationary Markov chain as the optimal fitting parameters.

### Nematodes’ trajectories in wind boundary layers

The motion of nematodes in the wind’s boundary layers is modeled by:

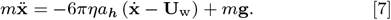

Here, **U**_w_ is the wind velocity, whose *x*- and *y*-components are modeled using the Blasius solution of a laminar boundary layer (49):

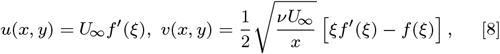

where *U*_*∞*_ is the free stream wind speed and *ν* = 1.562 × 10^*−*5^ m^2^/s is the kinematic viscosity of air, and 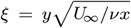 is a self-similar dimensionless variable. The function *f* (*ξ*) is the solution of the ordinary differential equation: 2*f* ^*′′′*^ + *f* ^*′′*^*f* = 0, where the prime denotes the differentiation with respect to *ξ*. An analytical approximation of *f* (*ξ*) was used and discussed in the *SI* (50). Our experimental setup using folded wet filter papers as a platform for the nematodes may induce discrepancies with the Blasius solution, which assumes a flat plane. However, this will not qualitatively affect our result, as the flow is predominantly along the long edge of the paper (see *SI* and Movie S3). We fitted the model in Eq. (7) and Eq. (8) to nematodes’ trajectories in two dimensions. The 2D model has a set of 5 fitting parameters: **Π** = [*a*_*h*_, *u*_0_, *v*_0_, *x*_0_, *U*_*∞*_], where *x*_0_ denotes the horizontal distance between the nematodes’ jumping location and the leading edge of the plate (i.e., *x* = 0). We then used the aforementioned MCMC method to fit our experimental data to the 2D model.

## ACKNOWLEDGMENTS

This work was supported by a grant from the W. M. Keck Foundation (to J.C.B. and R.R.), and the Tarbutton Postdoctoral Fellowship of Emory College of Arts and Sciences (to R.R.).

