## Supporting Information for "Electrostatics facilitate mid-air host attachment in parasitic jumping nematodes"

#### This PDF file includes:

Supporting text
Figs. S1 to S8
Table S1
Legends for Movies S1 to S7
SI References

#### Other supporting materials for this manuscript include the following:

Movies S1 to S7

### Supporting Information Text

### Experimental Setup

Figure S1($a$) shows the experimental setup for electrostatic experiments. Water drops containing living *Steinernema carpocapsae* nematodes were deposited on a vertically orientated wet filter paper sheet. The wet filter paper was folded back and forth to have a thickness of greater than 1 mm ($\approx 50$ times the nematode thickness) and was connected to the ground through a metal stand. A fruit fly was tethered to a copper wire and connected to a high-voltage power supply (ES5P-10W, Gamma High Voltage Research, Inc.). The voltage on the tethered fly was adjusted from $100-700$ V relative to the ground. The grounded paper was placed at two different heights below the fly, 5.1 mm and 6.2 mm, respectively. Analogous electrostatic experiments were performed by replacing the fruit fly by a charged metal sphere with a diameter of 2.54 mm. Figure S1($b$) shows the experimental setup for nematodes drifting in wind. Nematodes jumped from the same folded wet filter paper with air flows generated by a wind tunnel. The flow speed was approximately 0.2 m/s. In both electrostatic and wind experiments, nematodes with successful jumping were recorded using a high-speed camera (Nova S6, Photron USA, Inc.) at a frame rate of 10,000 frames/s.

### Takeoff and landing in still air

As a control, we recorded nematodes jumping in still air, without wind and electrostatic effects. Due to their submillimeter size, nematodes have a small Reynolds number ($Re \sim 10^0$) and experience significantly larger air drag forces compared to inertial forces. Consequently, their horizontal velocity during takeoff is greatly damped by air drag upon landing, resulting in an inclined takeoff and a vertical landing, as shown in Figs. S2($a$) and ($b$). This observation is consistent with our numerical simulations that model the nematodes as spheres (main text, Fig. 2$C$). However, a vertical landing is not always the case. Nematodes experience a drag force perpendicular to their elongated bodies. As a result, the landing angle can vary depending on their body orientation. Figure S2($c$) shows an example of a nematode with an inclined landing despite having a vertical takeoff, due to reorientation of its body. It has been suggested that *S. carpocapsae* have the ability of ariel righting (i.e., the ability to reorient themselves in midair), by maintaining a curved body posture to stop their rotations during landing (1). This stop of body rotation is observed in Figs. S2($a$), ($b$), and ($c$). Nonetheless, further research is needed to better understand whether nematodes can utilize their aerial righting ability to achieve directional landing.

### Trajectory Fitting in Three Dimensions

Since electrostatic forces depend on the three-dimensional (3D) distance between nematodes and hosts, we fit the experimental trajectories of jumping nematodes to our theoretical model in three dimensions. Figures S3($a$) and S4($a$) show the 3D fitting results and experimental data for nematodes' trajectories attracted by a fruit fly and by a charged metal ball, respectively. The two-dimensional (2D) projection of these 3D trajectories are shown in Figs. S3($b$) and S4($b$), respectively. For both the fruit fly and the metal ball data, the nematodes' trajectories exhibit considerable out-of-plane displacement and velocity components, indicating the necessity of 3D fitting. Figure S5($a$) shows the experimental data for a nematode's displacement, $x(t)$ and $y(t)$, as a function of time (red and blue symbols), and their model fitting results (solid black curves). Figure S5($b$) shows the 2D displacement in space, $\mathbf{x} = x\hat{\mathbf{x}} + y\hat{\mathbf{y}}$, for both experimental data (red dots) and model fitting (black curves). We notice that the model fitting results capture the nematode's displacement extraordinarily well, both as a function of time and in space. Figure S5($c$) shows the 3D displacement of the nematode, $\mathbf{x} = x\hat{\mathbf{x}} + y\hat{\mathbf{y}} + z\hat{\mathbf{z}}$, with the inferred the out-of-plane displacement, $z(t)$. This illustrates the difference between the full 3D displacement (black curves), the 2D displacement (blue curves), and the experimental data (red dots).

### Markov chain Monte Carlo method

Our theoretical model [Eq. (1), main text] has a set of 6 fitting parameters: $\mathbf{\Pi} = [q,\, a_h,\, u_0,\, v_0,\, w_0,\, z_0]$. Here, $q$ and $a_h$ are the charge and the hydrodynamic radius of a jumping nematode, $u_0$, $v_0$, $w_0$ are the three components of the nematode's initial or jumping velocity: $\mathbf{U}_0 = u_0\hat{\mathbf{x}} + v_0\hat{\mathbf{y}} + w_0\hat{\mathbf{z}}$, and $z_0$ is the out-of-plane component of the nematode's initial position. We want to find the optimal set of parameters $\mathbf{\Pi}_{\mathrm{M}}$ that can make predictions on data $\mathbf{X} = [x,\, y]$, where $x$ and $y$ are the experimental measurement of the nematode's position. Statistically, we would like to find the arguments that can maximize the conditional probability $P(\mathbf{\Pi}|\mathbf{X})$:

$$\mathbf{\Pi}_{\mathrm{M}} = \underset{\mathbf{\Pi}}{\operatorname{argmax}}\, P(\mathbf{\Pi}|\mathbf{X}). \tag{1}$$

Although an explicit expression for $P(\mathbf{\Pi}|\mathbf{X})$ is not available, its values can be computed using Bayes' theorem:

$$P(\mathbf{\Pi}|\mathbf{X}) = \frac{P(\mathbf{X}|\mathbf{\Pi})P(\mathbf{\Pi})}{P(\mathbf{X})}. \tag{2}$$

Here, $P(\mathbf{X}) = \int P(\mathbf{X}|\mathbf{\Pi})P(\mathbf{\Pi})\, d\mathbf{\Pi}$ is a normalizing constant that can be dropped (2). The prior probability $P(\mathbf{\Pi})$ is assumed to satisfy a multivariate normal distribution $\mathbf{\Pi} \sim \mathcal{N}(\mathbf{\Pi}_0, \sigma_0^2\mathbf{A})$, where $\mathbf{\Pi}_0$ is an initial guess, $\sigma_0$ is the uncertainty of the initial guess, and $\mathbf{A}$ is a diagonal covariance matrix such that each component of $\mathbf{\Pi}$ is assumed to be independent. The prior probability can be then estimated as:

$$P(\mathbf{\Pi}) = \left(1/\sqrt{2\pi\sigma_0^2}\right)^n \det(\mathbf{A})^{-1/2} \exp\left(-\,(\mathbf{\Pi} - \mathbf{\Pi}_0)^{\mathrm{T}}\mathbf{A}^{-1}(\mathbf{\Pi} - \mathbf{\Pi}_0)\,/2\sigma_0^2\right), \tag{3}$$

where $n = 6$ is the total number of dimensions for the six-parameter model. The conditional probability $P(\mathbf{X}|\mathbf{\Pi})$ is defined to minimize the loss function between experimental data and model prediction. We denote the model prediction on experimental data $\mathbf{X}$ using parameters $\mathbf{\Pi}$ as $\mathbf{X}'$, and the loss function is defined using $L^2$-norm as:

$$\mathcal{L}(\mathbf{X}, \mathbf{X}') = \left\|\mathbf{X} - \mathbf{X}'\right\| = \sqrt{\sum_{i=1}^{n}(x_i - x_i')^2 + (y_i - y_i')^2}, \quad [4]$$

where $x'$ and $y'$ denote the model's predictions on $x$ and $y$, and $x_i = x(t_i)$ is the measurement at a given discrete time $t_i$. The loss function is zero for an "ideal" model and we assume the loss function is normally distributed around zero: $\mathcal{L}(\mathbf{X}, \mathbf{X}') \sim \mathcal{N}(0, \sigma^2)$. The conditional probability can be then estimated as:

$$P(\mathbf{X}|\mathbf{\Pi}) = \left(1/\sqrt{2\pi\sigma^2}\right) \exp\left(-\mathcal{L}^2(\mathbf{X}, \mathbf{X}')/2\sigma^2\right). \quad [5]$$

With the evaluation of $P(\mathbf{\Pi})$ and $P(\mathbf{X}|\mathbf{\Pi})$ in Eq. (3) and Eq. (5), we then sampled from $P(\mathbf{\Pi}|\mathbf{X})$ using the Metropolis-Hastings algorithm (3, 4), which can generate a Markov chain with the following steps:

Step 0. Propose an initial guess $\mathbf{\Pi} = \mathbf{\Pi_0}$.

Step 1. Denote the current state as $\mathbf{\Pi}$. Propose a new state $\mathbf{\Pi'} = \mathbf{\Pi} + \mathbf{A}\boldsymbol{\delta}$ using a Gaussian random walk $\boldsymbol{\delta} \sim \mathcal{N}(\mathbf{0}, \sigma_{\mathrm{rw}}^2\mathbf{I})$, where $\sigma_{\mathrm{rw}}$ is the step of the random walk, and $\mathbf{0}$ and $\mathbf{I}$ are the zero vector and the identity matrix, respectively.

Step 2. Calculate the acceptance ratio $\alpha$:

$$\alpha = \min\left(1, \frac{P(\mathbf{X}|\mathbf{\Pi'})P(\mathbf{\Pi'})}{P(\mathbf{X}|\mathbf{\Pi})P(\mathbf{\Pi})}\right). \quad [6]$$

Step 3. Generate a random number $u$ from a uniform distribution $\mathcal{U}(0, 1)$. If $u < \alpha$, accept the new state $\mathbf{\Pi} = \mathbf{\Pi'}$; otherwise reject $\mathbf{\Pi'}$ and keep $\mathbf{\Pi} = \mathbf{\Pi}$. Return to Step 1.

We repeat the above sampling steps for 7,500 to 10,000 times until the loss function converges to its minimum. After the convergence, we continue the sampling for an additional 7,500 to 10,000 times to generate a stationary Markov chain, where all components of $\mathbf{\Pi}$ fluctuate around their mean values, as shown in Fig. S6. We then use the mean values of $\mathbf{\Pi}$ on the stationary Markov chain to approximate $\mathbf{\Pi}_{\mathrm{M}}$, the optimal fitting parameters.

### Justification of electrostatic model assumptions

In our model, we have assumed that the fruit fly host is an isolated charged sphere, and that the folded wet filter paper can be modeled as a grounded infinite plane. These assumptions simplify the model so that it is computationally feasible to calculate the best fit trajectory of a worm in an iterative procedure. But these assumptions may affect our inference of the nematode charge. The first assumption (isolated sphere) introduces a relatively small error. The relationship between the potential ($\phi$) and net charge ($Q$) on an isolated sphere of radius $a$ is $Q = 4\pi\epsilon_0 a\phi$. In the presence of a grounded plane a distance $h$ below the sphere, the capacitance of the system changes and the relationship can be written as:

$$Q = 4\pi\epsilon_0\phi\left(1 + \frac{a}{2h} + \mathcal{O}\left(\frac{a}{h}\right)^2\right), \quad [7]$$

where we have assumed $h > a$. In our experiments, $h \approx 6$ mm and $a \approx 1$ mm. Thus, the presence of the plane increases the charge on the host by about 10%, which is small compared to the spread of the inferred charge on the worm, $q$ (see Fig. 3*B*, main text).

The second assumption introduces a larger potential error. As stated, the folded, wet filter paper is approximately 1 mm thick, and was used in the experiments so that the nematodes mostly remained in the depth of field of the imaging system, and could tracked accurately. While 1 mm is much larger than the size of the nematodes, it is comparable to the size of the fruit fly host. In reality, the wet filter paper is more like a wedge with a rounded tip of radius $\sim 0.5$ mm. Charge tends to accumulate near sharp tips and edges, so we may expect that the surface charge (and thus $q$) would be larger in magnitude than the expectation from an infinite plane. To estimate the size of this effect, we consider just the rounded portion of the filter paper, and model the edge as an infinitely long, conducting cylinder of radius $R$. The fruit fly host is positioned at a distance $(R + h)$ above the axis of the cylinder. We are interested in the charge density ($\sigma$) on the surface of the cylinder immediately below the host. If we assume the charged host is a point charge, this electrostatic problem can be represented as an infinite sum (5):

$$\sigma = -\frac{Q}{2\pi R^2}\sum_{m=-\infty}^{\infty}\int_0^{\infty}\frac{K_m(k(R+h))}{K_m(kR)}dk, \quad [8]$$

where $K_m$ is the modified Bessel function of the second kind of order $m$. In the limit of a very large cylinder ($R \to \infty$), this sum reduces to the result for an infinite plane, $\sigma_p = Q/2\pi h^2$. For $h \approx 6$ mm and $R \approx 1$ mm, the charge density is enhanced by $\sigma/\sigma_p \sim 2.6$.

In reality, charge is not just distributed near the rounded edge of the filter paper, so the true enhancement is likely smaller than a factor of 2.6. However, this enhancement would increase the magnitude of the electric field, and a smaller value of $q$ would be necessary to fit the nematode trajectories. Nevertheless, the spread in the data in Fig. 3$B$ in the main text is quite large, and 0.1 pC would still be a reasonable estimate of the nematode charge. Ultimately we choose the simplest model for our computational inference procedure, and an error in the inference of $q$ would not change the conclusion that induction is the electrostatic mechanism that controls the charge on the nematodes.

### Numerical Simulations of the Charge on a Conducting Cylinder

We used COMSOL Multiphysics® to simulate the electrostatics of an upright conducting cylinder on a grounded plane, similar a standing nematode on a wet filter paper in our experiments. The electric potential, $\varphi$, is determined by Laplace's equation:

$$0 = \nabla^2 \varphi. \qquad [9]$$

The electric field, $\mathbf{E}$, is the gradient of the electric potential:

$$\mathbf{E} = -\nabla \varphi. \qquad [10]$$

Figure S7($a$) shows the domain of the 2D axisymmetric electrostatic simulation, where a sphere at a electric potential $\phi$ was located at $y = 6.8$ mm and an image sphere at a potential $-\phi$ is located at $y = -6.8$ mm. The electric potential on the two spheres is adjusted between $\phi = 100-700$ V in different simulations. A cylinder was placed between the two spheres with its bottom surface at $y = 0$. The height of the cylinder was fixed at 400 μm, similar to the length of a nematode. The radius of the cylinder varied between $a_1 = 20-200$ μm in different simulations using a parametric sweep. The symmetry of the two spheres makes $y = 0$ an effective grounded plane; the size of the cylinder is small enough compared to the two spheres so that the symmetry is not affected. To ensure grounding, the electric potential on the bottom surface of the cylinder was set to $\varphi = 0$. Figure S7($b$) shows the electric potential and the electric field lines in a simulation of $\phi = 500$ V and $a_1 = 20$ μm. We find that both the electric field and potential are highly symmetry about $y = 0$, except in the vicinity of the cylinder. The 3D illustration of the 2D axisymmetric electric potential is shown in Fig. S7($c$).

We calculated the electrostatic charge on the cylinder by integrating the normal component of the electric field over the cylinder's surface:

$$q = \frac{1}{\epsilon_0} \oiint_S (\mathbf{E} \cdot \mathbf{n}) \, \mathrm{d}A, \qquad [11]$$

where $\epsilon_0 = 8.854 \times 10^{-12}$ F/m is the vacuum permittivity. As a sanity check, we first simulated the charge on a conducting sphere of different radius $a$, and compared it to the theoretical prediction by Maxwell. Figure S8($a$) shows the simulation results of the charge on a sphere of $a = 100-200$ μm, along with Maxwell's prediction. We find that the simulation results agree extraordinarily well with theoretical prediction, validating our simulations. We proceeded with simulating the charge on a conducting cylinder of a fixed height of 400 μm and varying radius $a_1$. Figure S8($b$) shows the simulation results of the charge on a cylinder of $a_1 = 20-200$ μm. A comparison with Maxwell's prediction reveals that while the charge of a cylinder with $a_1 = 20$ μm is lower that of a sphere with $a = 100$ μm, the charge of a cylinder with $a_1 = 40-200$ μm is comparable to that of a sphere with $a = 100-200$ μm, similar to the inferred charge on nematodes in our experiments. This suggests that the observed nematode charge in our experiments (green shaded area, Fig. S8 and Fig. 3$B$, main text) is larger than the induced charge on a cylinder of the size of a worm. This difference can be attributed to the factor of 2.6 derived in the previous section, due to the geometry of the grounded wet filter paper.

### Numerical Simulations of nematodes drifting in wind with electrostatics

We performed numerical simulation of nematodes drifting in wind with a charged host nearby. In our simulations, the center of a charged sphere of a radius $a = 1$ mm was located at $x = 0$ and $y = 6$ mm. The electric potential on the sphere, $\phi$, was adjusted between $\phi = 0-800$ V, with a stepwise increment of 25 V. The charge on the sphere was calculated as: $Q = 4\pi\epsilon_0 a\phi$, where $\epsilon_0$ is the vacuum permittivity. A 40-mm long grounded plate was placed at $y = 0$ with its leading edge at $x = -20$ mm and trailing edge at $x = 20$ mm. In each simulation, nematodes took off from the grounded plate with random initial positions uniformly distributed between $x_0 \in [-20, 20]$ mm. The jumping speed of each nematode was fixed at $U_0 = 1$ m/s with random takeoff angles uniformly distributed between $\theta_0 \in [\pi/4, 3\pi/4]$, or between 45° and 135° with respective to the grounded plate. A horizontal laminar flow, $\mathbf{U}_\mathrm{w} = u\hat{\mathbf{x}} + v\hat{\mathbf{y}}$, was included in the simulations with its components modeled by the Blasius solution:

$$u(x, y) = U_\infty f'(\xi), \ \ v(x, y) = \frac{1}{2}\sqrt{\frac{\nu U_\infty}{x}} \left[\xi f'(\xi) - f(\xi)\right], \qquad [12]$$

where the wind speed $U_\infty$ was adjusted from $0-0.6$ m/s with a stepwise increment of 0.025 m/s, $\nu = 1.562 \times 10^{-5}$ m²/s is the kinematic viscosity of air, and $\xi = y\sqrt{U_\infty/\nu x}$ is a self-similar dimensionless variable. The function $f(\xi)$ is the solution of the ordinary differential equation:

$$2f''' + f''f = 0, \qquad [13]$$

where the prime denotes differentiation with respect to $\xi$. The equation of motion for the jumping worm with both wind and electrostatics is now given by:

$$m\ddot{\mathbf{x}} = \frac{-Qq\left(\mathbf{x}-\mathbf{x}_0^+\right)}{4\pi\epsilon_0\left|\mathbf{x}-\mathbf{x}_0^+\right|^3} + \frac{Qq\left(\mathbf{x}-\mathbf{x}_0^-\right)}{4\pi\epsilon_0\left|\mathbf{x}-\mathbf{x}_0^-\right|^3} - 6\pi\eta a_h\left(\dot{\mathbf{x}}-\mathbf{U}_{\mathrm{w}}\right) + m\mathbf{g}. \qquad [14]$$

In each simulation, numerical integration of Eq. (14) was performed in MATLAB® for 1,000 hypothetical nematodes using the built-in function `ode45`. Each nematode has an identical hydrodynamic radius, $a_h = 100$ µm. The integration time for each nematode's trajectory is 0.5 s. Capture rate is calculated from 1,000 trajectories in each simulation, with "capture" defined as successful attachment to the charged sphere at $x = 0$ and $y = 6$ mm. To reduce the computation cost, we employed an analytical first-order approximation of $f(\xi)$, which is otherwise available only through numerical method (6):

$$f_1(\xi) = \xi - 1.8760 + 1.4564\mathrm{e}^{-\xi} + 1.2956\xi\mathrm{e}^{-\xi} + 0.4196\mathrm{e}^{-2\xi}. \qquad [15]$$

We compared the values of $f_1(\xi)$ and its first derivative $f_1'(\xi)$, to the exact value of $f(\xi)$ and $f'(\xi)$ from numerical method in the Table S1. We find the relative error is within 5% for $f_1(\xi)$ at $\xi > 2$ and within 7% for $f_1'(\xi)$ at $\xi > 1$, validating the analytic approximation.

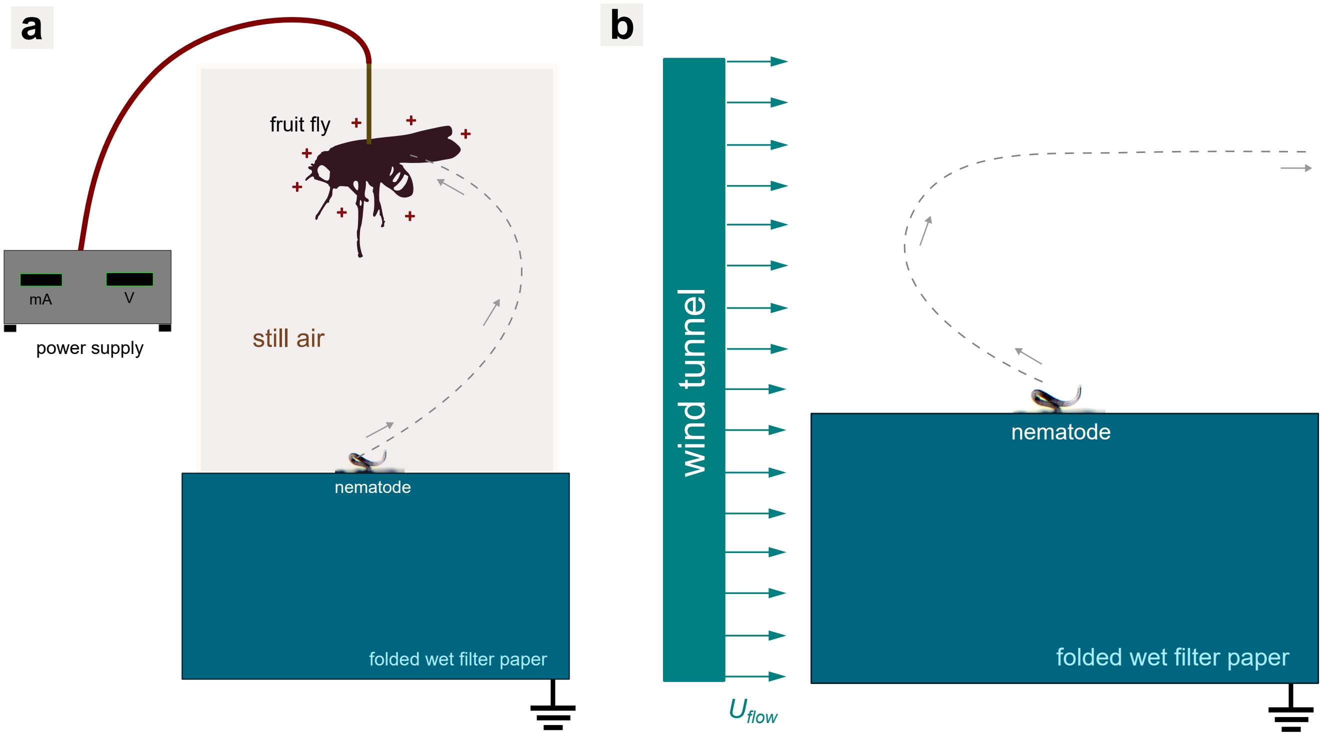


**Fig. S1.** Schematics of the experimental setup. (*a*) Host attachment of jumping nematodes with electrostatics. Nematodes launched from a vertically oriented sheet of folded wet filter paper to attach to their host, a charged fruit fly, which was tethered to a copper wire and connected to a high-voltage power supply. The wet filter paper was folded back and forth multiple time to have a certain thickness to stand, and was connected to the ground through a metal stand. (*b*) Ariel drifting of jumping nematodes in the wind. Nematodes took off from the same folded wet filter paper without a charged host nearby, and with a wind tunnel generating air flows.

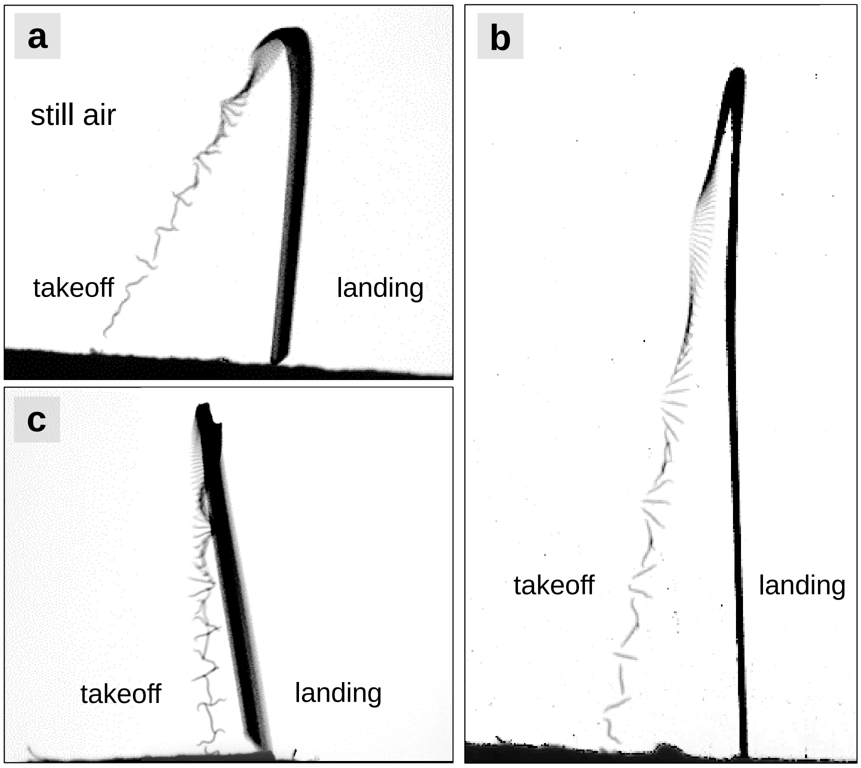


**Fig. S2.** Takeoff and landing of nematodes jumping in still air. (*a*) A nematode with an inclined takeoff and a vertical landing. (*b*) Another nematode with a inclined takeoff and a vertical landing. (*c*) A nematode with an vertical takeoff and a inclined landing due to body orientation.

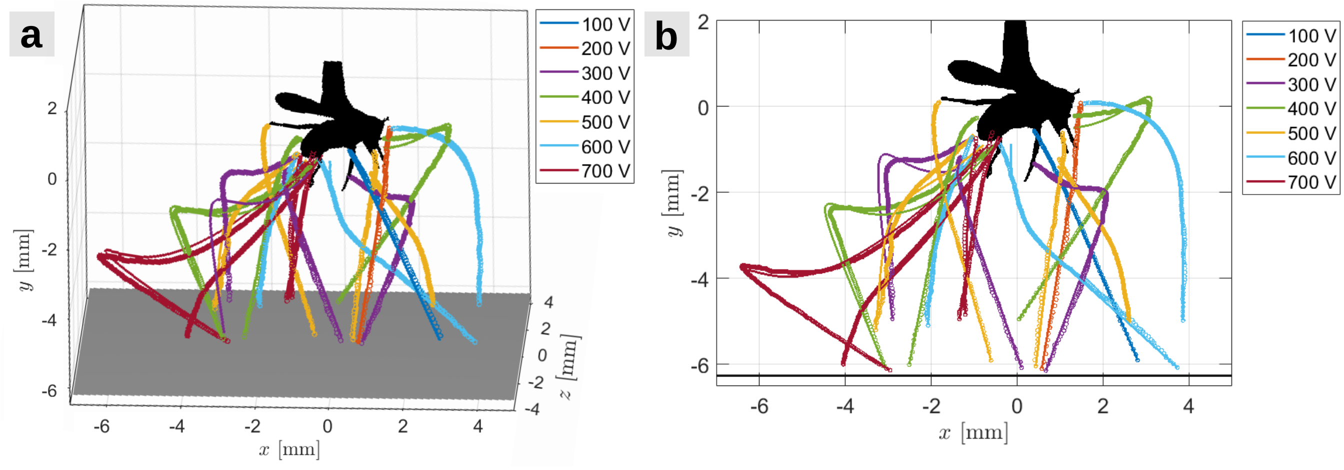


**Fig. S3.** Jumping nematodes electrostatically attracted by a charged insect host, a fruit fly. (*a*) Three-dimensional (3D) trajectories of the center of mass of jumping nematodes. (*b*) Two-dimensional (2D) projection of the 3D trajectories of jumping nematodes' center of mass. Circle symbols are experimental data, solid curves are model fitting results. Different colors represent different host's electric potentials.

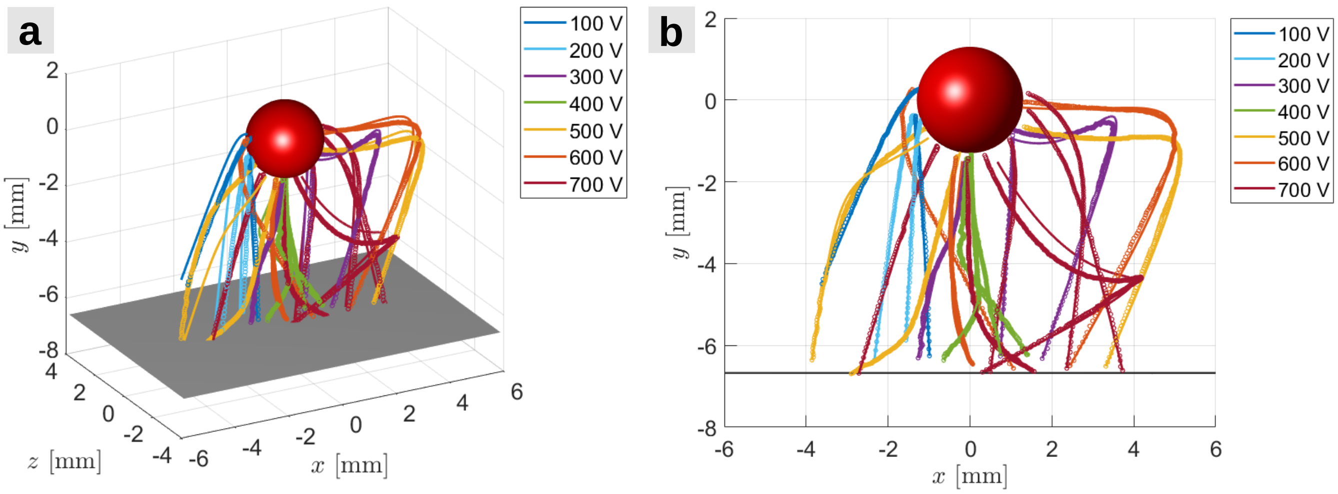


**Fig. S4.** Jumping nematodes electrostatically attracted by an analogous insect host—a charged metal sphere. (*a*) 3D trajectories of the center of mass of jumping nematodes. (*b*) 2D projection of the 3D trajectories of jumping nematodes' center of mass. Circle symbols are experimental data, solid curves are model fitting results. Different colors represent different host's electric potentials.

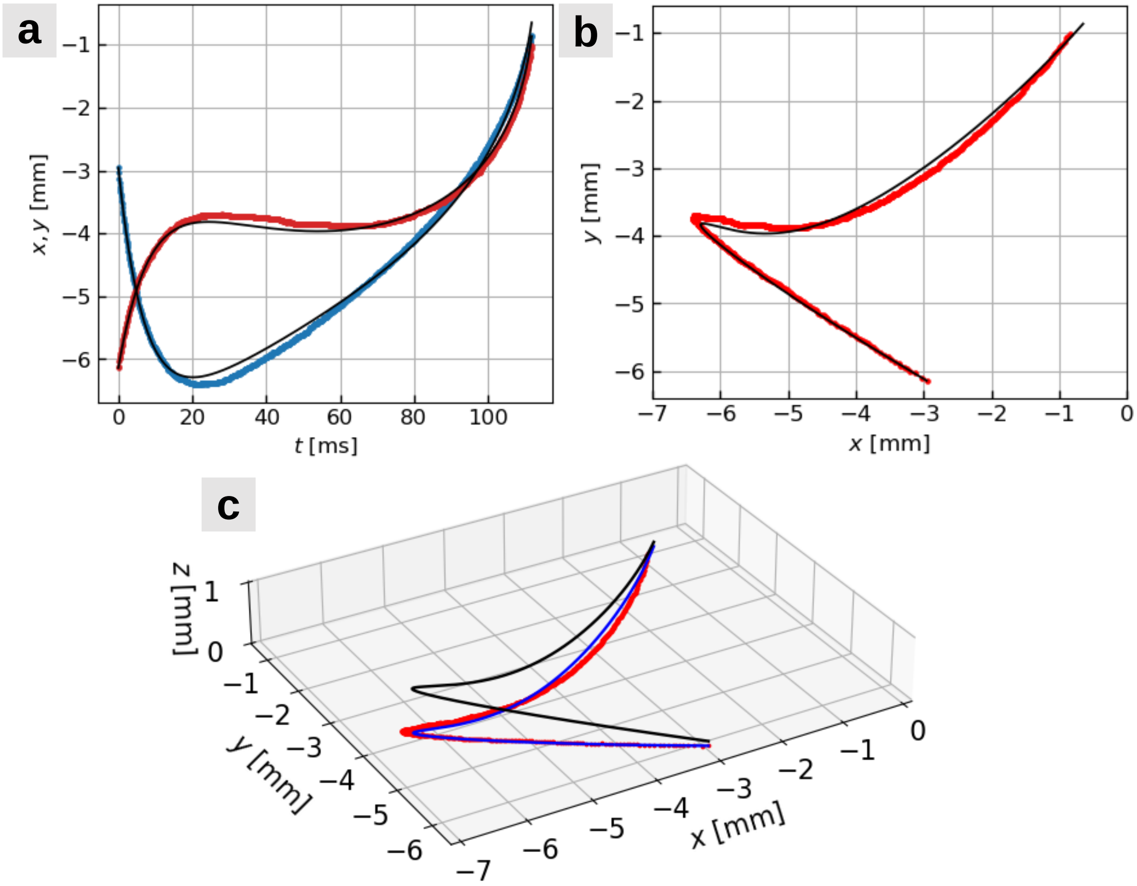


**Fig. S5.** Model fitting results of a nematode's trajectories with the voltage on the fruit fly at 700 V. (*a*) Experimental data (dots) and model fitting (black curves) for the jumping nematodes' $x$-position (red) and $y$-position (blue) as a function of time $t$. (*b*) Experimental data (red dots) and model fitting (black curves) for the trajectory of the jumping nematode in two dimensions. (*c*) Model fitted 3D trajectory of the jumping nematode (black curves), 2D projection of the 3D trajectory (blue curves), and comparison with the 2D experimental data (red dots).

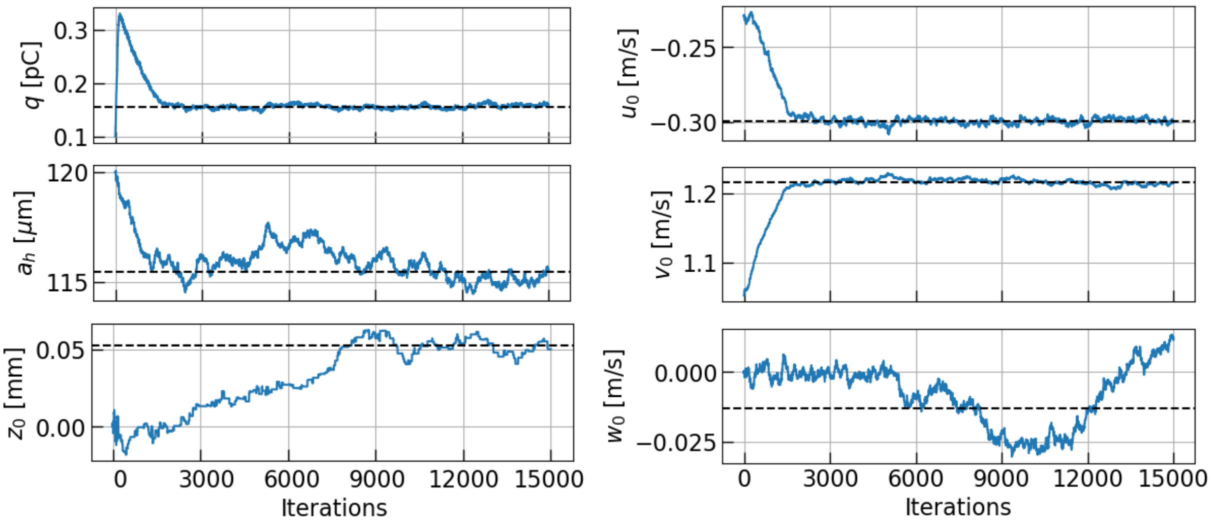


**Fig. S6.** Evolution of the fitting parameters: $q$, $a_h$, $z_0$, $u_0$, $v_0$, $w_0$, as a function of iteration number in a Markov chain. The inferred charged $q$, hydrodynamic radius $a_h$, in-plane velocities $u_0$ and $v_0$ become stationary in the Markov chain after around 3000 iterations, while the out-of-plane displacement and velocity $z_0$ and $u_0$ have more uncertainty and take more iterations to reach a stationary state. The black dashed lines represent the mean values of parameters in the stationary Markov chain, taken to be the optimal fitting parameters.

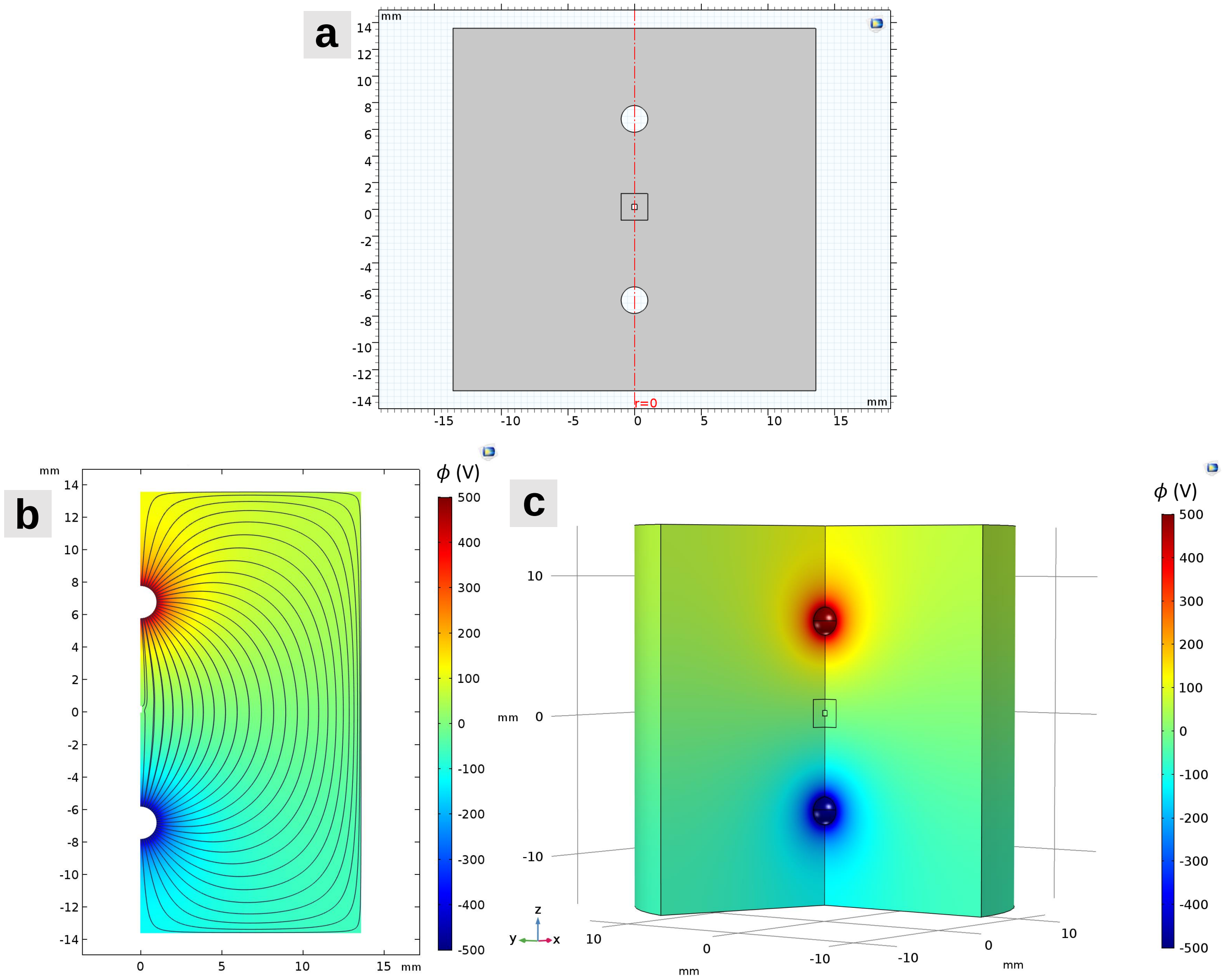


**Fig. S7.** Numerical simulations of the charge on an upright cylinder using the finite element method in COMSOL Multiphysics®. (*a*) Domain of the 2D axisymmetric electrostatic simulation. Two circles on the top and the bottom are the positively charged sphere and the negatively charged image sphere. A rectangle in the middle is a cylinder of fixed height of 400 $\mu$m and a varying radius $a_1$. The red dashed dotted line in the middle is the axis of symmetry. (*b*) 2D representation of the electric potential and electric field in numerical simulations. Color map represents the electric potential, and black curves are the electric field lines. The voltages on the top and bottom spheres were set to 500 V and $-500$ V, respectively. (*c*) 3D representation of the same electric potential data presented in (*b*).

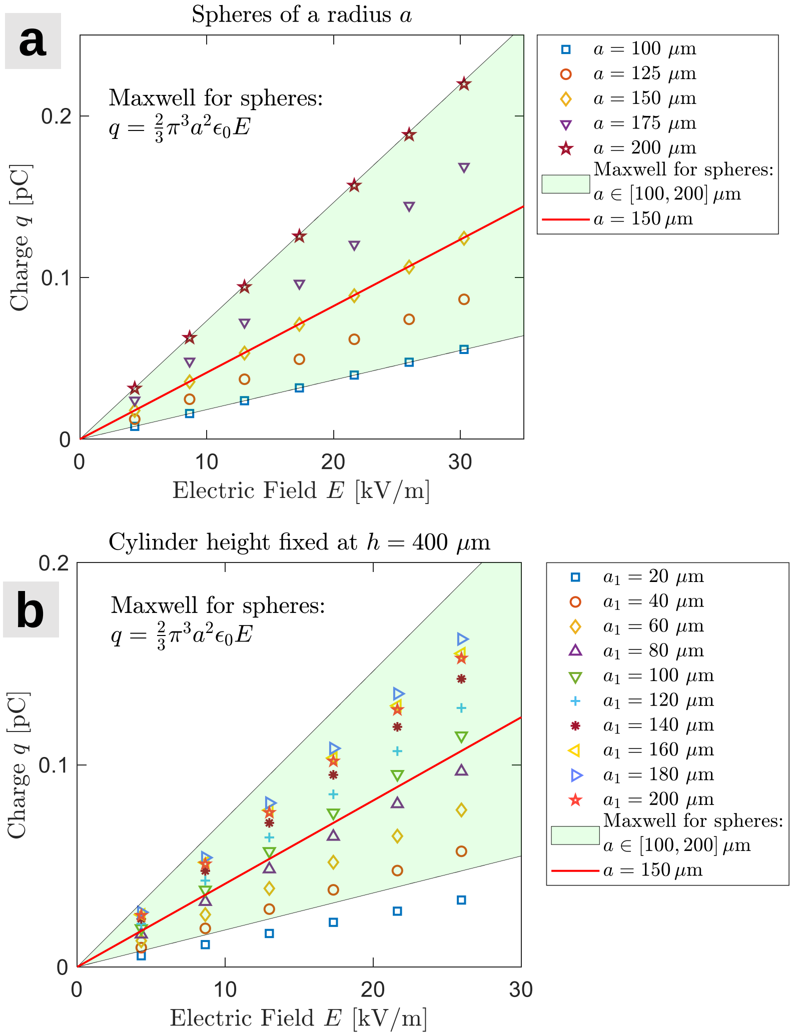


**Fig. S8.** The induction charge on spheres and cylinders from numerical simulations and comparison with Maxwell's prediction for spheres. (*a*) The induction charge on spheres of a radius $a = 100-200$ µm (markers) and Maxwell's prediction for spheres (shaded area and red line). The numerical data agree almost perfectly with theoretical predictions. (*b*) The induction charge on upright cylinders of a fixed height of 400 µm and varying radius of $a_1 = 20-200$ µm (markers) and Maxwell's prediction for spheres (shaded area). The numerical data for cylinders of radius $a_1 = 40-200$ µm lie within with theoretical predictions for spheres of a radius $a = 100-200$ µm.

**Table S1. Comparison between the exact values of the Blasius function $f(\xi)$ and its derivative $f'(\xi)$ from numerical method, and the analytical approximation $f_1(\xi)$ and $f_1'(\xi)$.**

| $\xi$ | $f(\xi)$ | $f_1(\xi)$ | error | $f'(\xi)$ | $f_1'(\xi)$ | error |
|---|---|---|---|---|---|---|
| 0 | 0 | 0 | 0 | 0 | 0 | 0 |
| 0.5 | 0.04149 | 0.05462 | 31.66% | 0.16589 | 0.20083 | 21.07% |
| 1.0 | 0.16557 | 0.19319 | 16.68% | 0.32979 | 0.35065 | 6.32% |
| 1.5 | 0.37014 | 0.40349 | 9.01% | 0.48679 | 0.48871 | 0.39% |
| 2.0 | 0.65003 | 0.67947 | 4.53% | 0.62977 | 0.61219 | 2.79% |
| 2.5 | 0.99631 | 1.01225 | 1.60% | 0.75126 | 0.71527 | 4.79% |
| 3.0 | 1.39682 | 1.39106 | 0.41% | 0.84605 | 0.79640 | 5.87% |
| 3.5 | 1.83770 | 1.80530 | 1.76% | 0.91304 | 0.85745 | 6.09% |
| 4.0 | 2.30576 | 2.24573 | 2.60% | 0.95552 | 0.90185 | 5.62% |
| 4.5 | 2.79013 | 2.70500 | 3.05% | 0.97951 | 0.93334 | 4.71% |
| 5.0 | 3.28329 | 3.17748 | 3.22% | 0.99155 | 0.95523 | 3.66% |
| 5.5 | 3.78057 | 3.65908 | 3.21% | 0.99688 | 0.97021 | 2.68% |
| 6.0 | 4.27964 | 4.14688 | 3.10% | 0.99898 | 0.98033 | 1.87% |
| 7.0 | 5.27926 | 5.13360 | 2.76% | 0.99992 | 0.99158 | 0.83% |
| 8.0 | 6.27923 | 6.12796 | 2.41% | 1.00000 | 0.99618 | 0.18% |

**Movie S1. Slow motion video of a jumping nematode attracted by a charged insect host (a fruit fly). The video plays at 0.012 times the real-time speed.**

**Movie S2. Slow motion video of a Jumping nematode attracted by a charged metal sphere, a hypothetical host. The video plays at 0.012 times the real-time speed.**

**Movie S3. Slow motion video of a jumping nematode drifting in a horizontal laminar flow generated by a wind tunnel. The video plays at 0.012 times the real-time speed.**

**Movie S4. Numerical simulations of nematodes drifting in wind with a charged spherical host nearby. The host's electric potential was increased from $\phi = 0$ to $\phi = 800$ V, at several fixed wind speeds, ranging from $U_\infty = 0 - 0.6$ m/s.**

**Movie S5. Numerical simulations of nematodes drifting in wind with a charged spherical host nearby. The wind speed was increase from $U_\infty = 0$ to $U_\infty = 0.6$ m/s, at several fixed host's electric potentials, ranging from $\phi = 0 - 800$ V.**

**Movie S6. Slow motion video of a standing nematode attracted by a plastic syringe rubbed on human hair. The video plays at 0.003 times the real-time speed.**

**Movie S7. Slow motion video of a a group of nematodes attracted by a charged water droplet. The video plays at 0.003 times the real-time speed.**
